# Gene expression patterns in chordate embryonic development suggest partial applicability of Haeckel’s postulates

**DOI:** 10.1101/2020.08.25.265843

**Authors:** Song Guo, Haiyang Hu, Chuan Xu, Naoki Irie, Philipp Khaitovich

## Abstract

The relationship between embryonic development and evolution historically investigated based on embryo morphology, could now be reassessed using mRNA expression endophenotype. Here, we investigated the applicability of von Baer’s and Haeckel’s arguments at mRNA expression level by comparing the developmental changes among nine evolutionarily distinct species: from oyster to mouse. In agreement with models based on von Baer’s postulates, up to 36% of mRNA expression indicated nearly linear conservation of species’ developmental programs. By contrast, 5-15% of developmental expression profiles, enriched in neural genes, displayed an alignment pattern compatible with the terminal edition paradigm proposed by Haeckel. Thus, the development-evolution relationship based on mRNA expression agrees with early concepts based on embryo morphology and demonstrates that the corresponding patterns coexist in chordate development.

## Introduction

The general relationship between ontogeny and phylogeny, or development and evolution, has long been discussed. Among cornerstones of the evolutionary developmental theory are the arguments formulated in von Baer’s laws of embryology and Ernst Haeckel’s Biogenetic law. Although von Baer did not accept the concept of evolution, his idea that the earlier stages of embryogenesis reflect shared traits of a broader taxonomic group(Von Baer, 1828) laid the foundation for the current evolutionary views on developmental programs’ conservation. Ernst Haeckel, who advocated evolution, proposed in his Biogenetic law that the embryogenesis replays the species’ evolutionary past. Thus, according to this concept, the new developmental stages will be added to the ancestral embryonic program to produce more recently evolved species(Haeckel, 1866). After a long debate, the general concept of early embryogenesis conservation, reflecting a more ancestral stage, continued in a form of a monotonic developmental conservation model known as the funnel model. More recently, a developmental conservation model called the developmental hourglass model was proposed, which placed the most significant stage conservation at a mid-embryonic part of embryogenesis, defined as the phylotypic period(Duboule, 1994). The applicability of the hourglass model to the conservation of developmental stages at mRNA expression level has been supported both in animals(Domazet-Loso & Tautz, 2010; Hu et al., 2017; Irie & Kuratani, 2011; Irie & Sehara-Fujisawa, 2007; Kalinka et al., 2010) and plants(Cheng, Hui, Lee, Wan Law, & Kwan, 2015; Cridge, Dearden, & Brownfield, 2016; Quint et al., 2012). Recent studies, however, demonstrated the applicability of the funnel model or the coexistence and validity of both funnel and hourglass models at different levels of trait evolution in vertebrates(Artieri, Haerty, & Singh, 2009; Bininda-Emonds, Jeffery, & Richardson, 2003; Comte, Roux, & Robinson-Rechavi, 2010; Hu et al., 2017; Levin et al., 2016; Piasecka, Lichocki, Moretti, Bergmann, & Robinson-Rechavi, 2013; Roux & Robinson-Rechavi, 2008; Uesaka, Kuratani, Takeda, & Irie, 2019). Even the broadly criticized Biogenetic law, has been pointed out to have potential applicability of its principles(Richardson & Keuck, 2002).

One of the major barriers in testing the relationship between development and evolution, including the applicability of concepts proposed in Biogenetic law or the von Baer’s law of embryology, is the difficulty in identifying evolutionarily-homologous developmental stages among distant species. While pioneering studies focused on the investigation of embryonic morphology(Jeffery, Bininda-Emonds, Coates, & Richardson, 2002), recent works relied on developmental changes in mRNA expression profiles as an informative endophenotype(Bozinovic, Sit, Hinton, & Oleksiak, 2011; J. J. Li, Huang, Bickel, & Brenner, 2014; Piasecka et al., 2013; Yanai, Peshkin, Jorgensen, & Kirschner, 2011). The use of gene expression facilitates inter-species comparisons, as orthologous genes can be matched among species, and their expression profiles could be traced at all stages of development. Several studies examined developmental gene expression patterns in mammals(Wagner, Tabibiazar, Liao, & Quertermous, 2005), vertebrates(Piasecka et al., 2013), chordates(Levin et al., 2016; Yanai et al., 2011), and fruit fly species(Tomancak et al., 2007). These studies, however, either focused on gene expression patterns within a specific species or compared developmental stage conservation within the concepts of the hourglass model paradigm.

Here, we directly tested the compatibility of developmental expression patterns with predictions of the Biogenetic law and von Baer’s law of embryology. To do so, we performed temporal alignment of mRNA developmental expression trajectories among eight chordate species and oyster. Haeckel’s Biogenetic law proposing the addition of new parts to the ancestral developmental program (terminal addition) could be described as a “progressive” developmental model concerning the ontogenetic gene expression pattern alignment. By contrast, Baer’s law and its later modifications presume the existence of a general developmental program spanning the entire development, while more conserved at early embryonic stages. Such a model could be termed as a “continuous” developmental model concerning the ontogenetic gene expression pattern alignment.

## Results

### Alignment of developmental gene expression patterns among species

We investigated the relationship between ontogeny and phylogeny of chordate species at the level of molecular phenotype using RNA-sequencing (RNA-seq) data collected over the entire course of embryonic development in eight chordate species of different organizational complexity (amphioxus, ciona, zebrafish, two species of frogs, turtle, chicken, and mouse) and an outgroup species (oyster)(Hu et al., 2017; Zhang et al., 2012). In each species, the data were collected at 11 to 20 developmental stages and measured in duplicates (Figure 1A; Supplementary file 1: Table S1 and Figure 1A–figure supplement 1).

**Figure 1.**
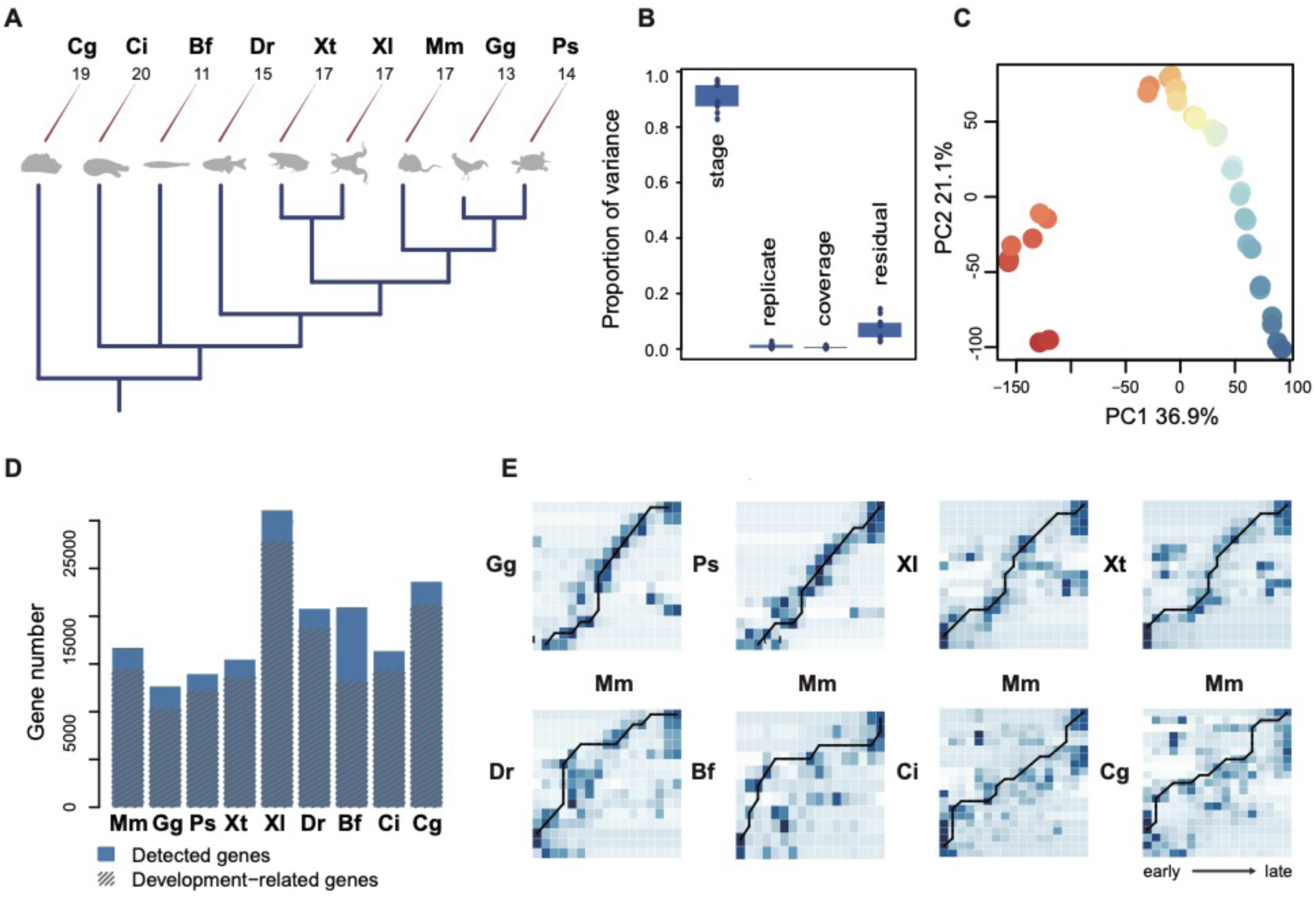
Species phylogeny and alignment of embryonic development stages. **A** Experimental scheme showing the phylogenetic relationship among species (dark blue dendrogram) and numbers of sampled developmental stages. The abbreviations here and throughout the figures indicate species: Cg – oyster (*Crassostrea gigas*), Ci – ciona (*Ciona intestinalis*), Bf – amphioxus (*Branchiostoma floridae*), Dr – zebrafish (*Danio rerio*), Xt – Western clawed frog (*Xenopus tropicalis*), Xl – African clawed frog (*Xenopus laevis*), Ps – turtle (*Pelodiscus sinensis*), Gg – chicken (*Gallus gallus*), Mm – mouse (*Mus musculus*). **B** Variance explained by different factors based on ANOVA. Boxes represent the interquartile values of the variance proportion explained by the factors listed on the x-axis among nine species. Dots represent the mean explained variance proportion for each species. **C** PCA plot based on the expression of 16,685 genes in mouse development. Dots represent samples, and color represents developmental stages (red – early, blue – late). **D** The number of detected and development-related genes in each species. **E** Heatmap showing the pairing scores reflecting overlap of stage-associate genes (darker shade of blue representing greater overlap) and the optimal alignment path calculated using the Needleman-Wunsch algorithm (black line).

Within species, differences among developmental stages explained approximately 90% of total expression variation (Figure 1B,C and Figure 1C–figure supplement 2). By contrast, other factors, such as biological replicates and sequence coverage, explained 1% and 0.6% of variation each (Figure 1B). Accordingly, on average, 85% of detected genes showed significant expression differences among developmental stages in each species (development-related genes) (polynomial test, Benjamini-Hochberg (BH) corrected *p* < 0.05; Figure 1D and Figure 1D-Source data 1).

We searched for the best temporal alignment between each pair of species using genes preferentially expressed at a particular developmental stage (stage-associated genes; Methods; Supplementary file 2: Table S2)(J. J. Li et al., 2014). Surprisingly, despite substantial differences in organizational complexity, an approximately linear alignment of developmental stages was a dominant stage-matching pattern in each pair of species (Figure 1E and Figure 1E–figure supplement 3,4).

### Two alternative evolutionary models of species’ development

We next sought to test, from the perspective of gene expression, two models, the progressive and continuous one, describing embryogenesis rooted in von Baer’s and Haeckel’s ideas. Specifically, we based the progressive model (PM) on Haeckel’s postulate that embryogenesis replays the evolutionary history of the species. Based on this view, the developmental gene expression profiles of a species with more ancestral body plan organization will align the best to early embryonic stages of species with more recently evolved organization, “replaying” the evolutionary history of this body plan (Figure 2A; Figure 2A–figure supplement 5). By contrast, evolutionary developmental models rooted in von Baer’s ideas, such as the hourglass and the funnel model, presume general preservation of the developmental profiles across species (Figure 2A). Accordingly, we based an alternative continuous model (CM) on this assumption, thus predicting the best alignment between complete, non-truncated, developmental expression profiles of the species.

**Figure 2.**
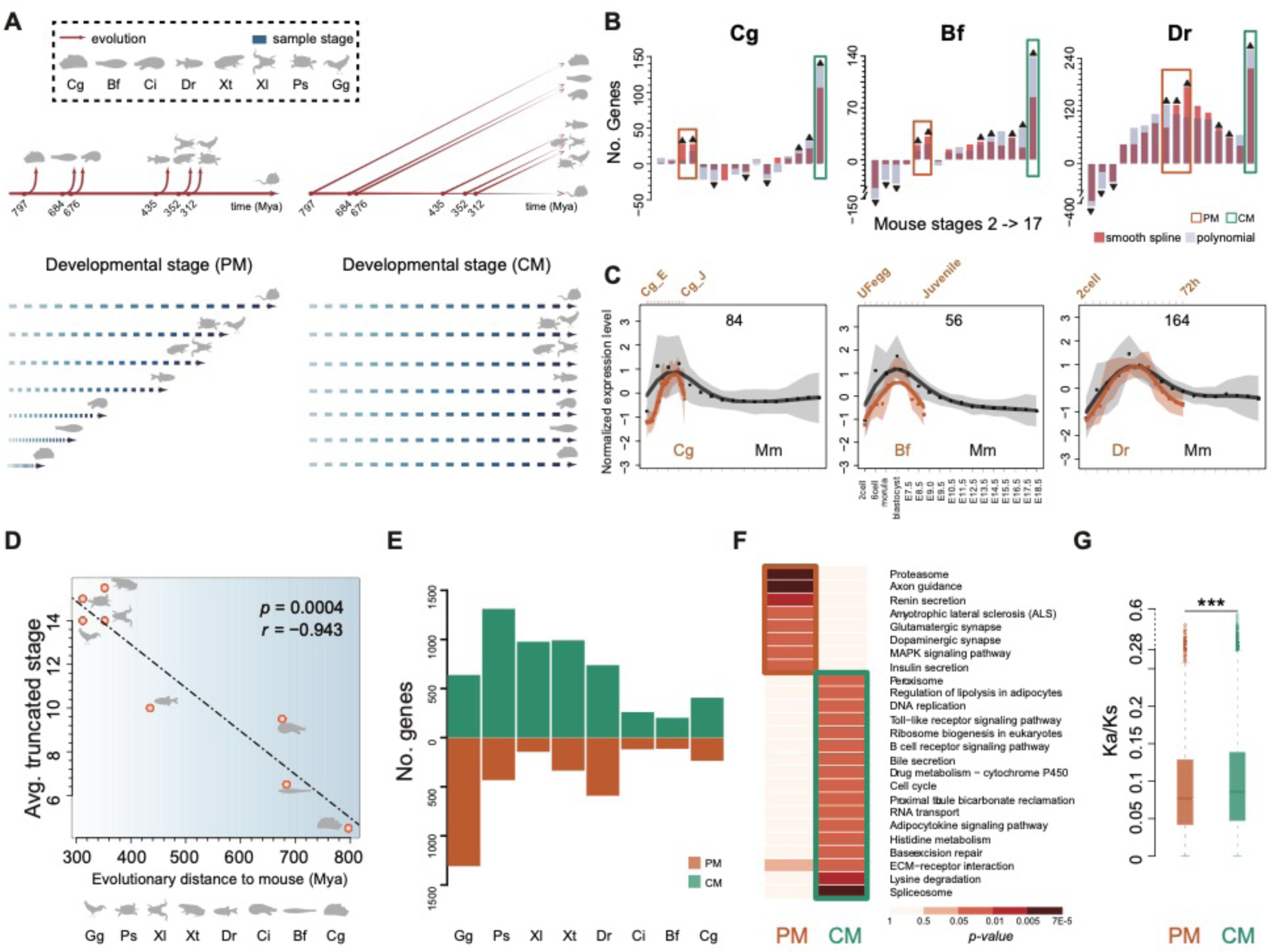
Relationship of developmental gene expression patterns among species. **A** Schematic description of the progressive and continuous models and their predicted alignment patterns. The progressive model (PM, left) involves an extension of developmental program in more recently evolved species, thus predicting the best alignment to a shortened mouse developmental course. The continuous model (CM, right) predicts the best alignment to the complete developmental course. **B** Numbers of genes showing the best expression trajectory’s alignment between the complete developmental course of the oyster (left), amphioxus (middle), and zebrafish (right) and mouse developmental sets of different lengths: from two to 17 stages. Colors indicate two methods used for developmental expression trajectory calculation: cubic smooth spline (orange) and polynomial regression (gray). Colored rectangles mark stages containing alignments fitting CM (green) or PM (orange) predictions. Black triangles indicate the significantly greater (up) or lesser (down) number of genes than that expected by chance aligning to an indicated mouse developmental interval (permutation test, BH-corrected *p* < 0.05). **C** Examples of PM gene expression patterns. Dots show the cluster-level standardized expression levels at each developmental stage in mouse (black) and the other species (orange). The curves represent average expression profiles, and the shaded regions represent the standard deviation of curve estimates. **D** The relationship between the lengths of truncated sets of mouse stages containing maximal numbers of PM genes for each of the eight species and the phylogenetic distances to the mouse. Dots represent species. The dotted line marks the regression curve. The Pearson correlation coefficient and linear regression p-value are shown at the top right. Mya – millions of years ago. **E** Numbers of CM and PM genes identified in each of the eight species. Note that PM gene number in chicken (Gg) might be inflated due to poor resolution of PM and CM predictions at close phylogenetic distances. **F** KEGG pathways significantly enriched in PM or CM from all eight non-mouse species. The heat map shows the p-values of the enrichment tests. Colored rectangles mark significantly enriched KEGG pathways. **G** Amino acid sequence conservation (Ka/Ks) of PM and CM genes from all eight non-mouse species. Asterisks indicate the significance of the difference (Wilcoxon rank-sum one-sided test, *p* < 0.0005).

We found that out of all development-related genes that aligned significantly to the mouse developmental trajectory, 19-36% aligned linearly, thus behaving in agreement with the continuous model predictions (CM genes) (correlation test, r > 0.7, *p* < 0.05; Figure 2B,E; Figure 2B–figure supplement 6 and Figure 2E – source data 1). By contrast, 5-15% of development-related genes aligned significantly better to the truncated sets (PM genes) (permutations, BH-corrected *p* < 0.05; Figure 2B,C,E; Figure 2B–figure supplement 6 and Figure 2E – source data 2). Further, in each pairwise alignment, PM genes did not align uniformly to all shortened mouse developmental intervals but tended to peak at a particular fragment. For instance, oyster PM genes aligned best to the mouse developmental fragments ending at stages 3-4, while amphioxus PM genes – to the fragments ending at stages 5-6 (Figure 2B,C and Figure 2C–figure supplement 7). Overall, PM genes in species evolutionarily proximal to the mouse tended to align to increasingly longer truncated mouse developmental sets (Figure 2D). This phylogenetic ordering of ontogenetic patterns was not caused by the alignment procedure, which was not biased to any developmental stage, and robust to the use of evolutionarily old genes present in each of the nine investigated species (Supplementary file 3: Figure S1). At the same time, this phenomenon matches the phylogeny-ontogeny relationship among multiple species proposed by Haeckel, even though we did not include this aspect in PM model formulation. By contrast, repeating the alignment procedure using species with more ancient body plans, such as amphioxus or ciona, instead of the mouse, did not yield a consistent significant excess of genes following PM predictions (Supplementary file 4: Figure S2).

### Properties of CM and PM genes

While we defined CM and PM gene sets independently for each species, each set overlapped significantly between species (Methods, subsampling, Bonferroni-corrected *p* < 0.05; Supplementary file 5: Table S3), indicating the conservation of CM and PM patterning across chordate embryogenesis and presumably convergent functionality specific to each model.

Indeed, CM genes showed enrichment in Gene Ontology (GO) terms corresponding to general cellular functions, such as spliceosome, RNA transport, DNA replication, and several metabolic processes (Hypergeometric test, *p* < 0.05; Figure 2F – source data 1). By contrast, PM genes were primarily enriched in neural functions, including axon guidance, glutamatergic synapse, dopaminergic synapse, and MAPK signaling pathways (Hypergeometric test, *p* < 0.05; Figure 2F and Figure 2F – source data 1). In line with functional enrichment results, expression of CM genes displayed significantly lower tissue-specificity in mouse(B. Li et al., 2017) compared to PM genes, and all expressed genes (one-sided Fisher’s exact test, *p* < 0.05; Supplementary file 6: Figure S3). Nonetheless, PM genes were more conserved at the amino acid sequence level compared to CM genes (Ka/Ks, one-sided Wilcoxon rank-sum test, *p* < 0.0005) (Figure 2G and Figure 2G–figure supplement 8,9).

### Visualization of developmental expression patterns

We next investigated developmental expression trajectories of the 244 orthologous genes classified as development-related in all nine species using the nearly linear, species’ alignment (Figure 3A). These genes were grouped into six clusters based on their developmental profiles in the unsupervised clustering analysis (Figure 3B). Remarkably, 66% of the genes fell within a single cluster representing a decreasing expression pattern conserved across all nine species (CL9.1) (Figure 3C). By contrast, the expression trajectories in the other clusters differed more among species, with the extent of differences being directly proportional to corresponding phylogenetic distances (Figure 3E and Figure 3E–figure supplement 10). Accordingly, CL9.1 genes substantially overlapped with CM genes (one-sided Fisher’s exact test, odds ratio = 3, *p* < 0.0001) and were shown in the same biological processes as CM genes, including spliceosome and RNA processing (Supplementary file 7: Table S4).

**Figure 3.**
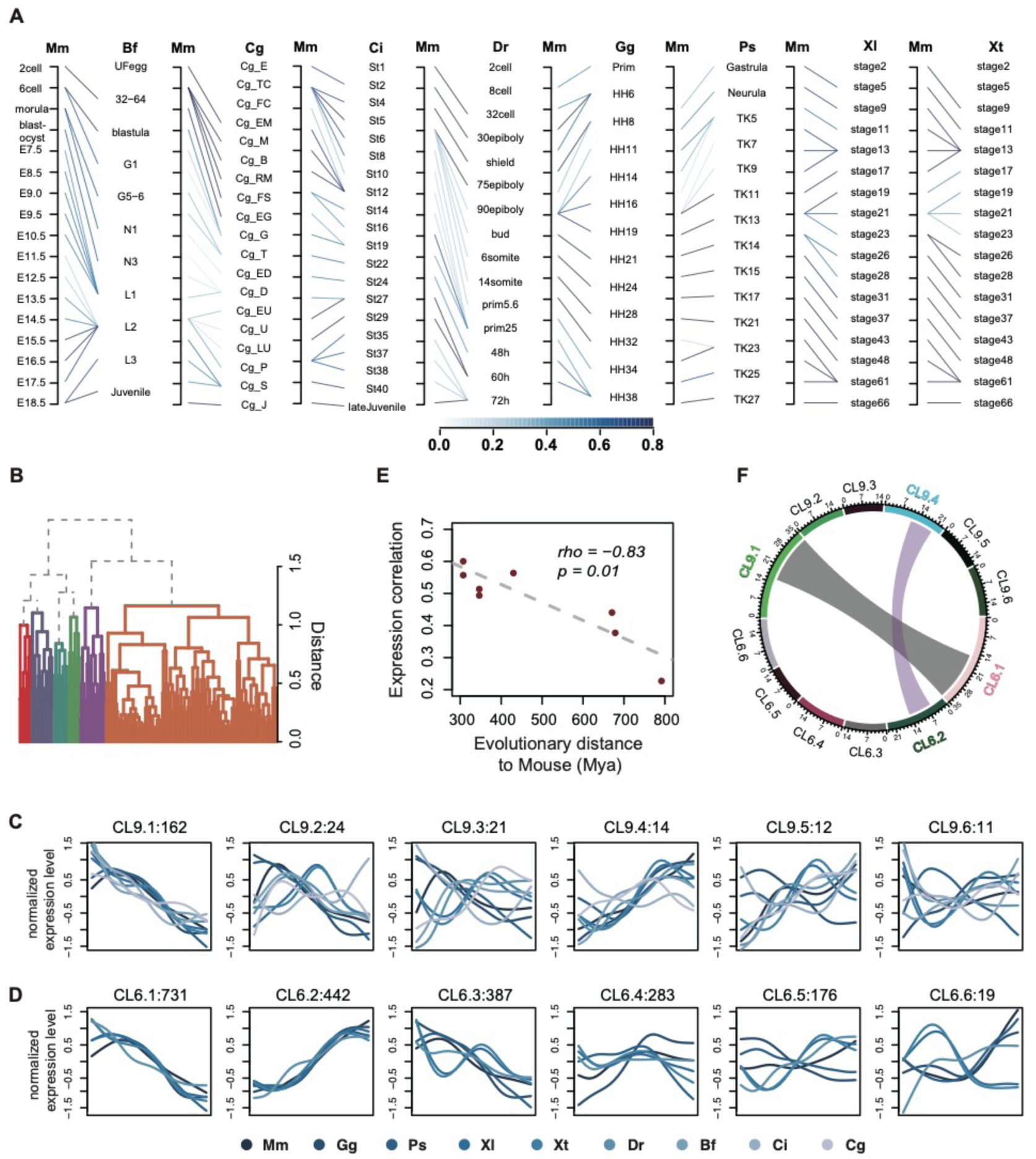
Developmental expression of genes based on the species’ alignment. **A** Pairwise alignment of developmental stages between mouse and the other species. Thick lines represent alignments supported by technical replicates. Thicker lines represent more stable alignments calculated by random subsampling of samples 500 times. **B** Hierarchical clustering of concatenated developmental gene expression trajectories of 244 gene orthologs shared among nine species. Colors represent clusters. **C** Developmental gene expression patterns in each of six clusters based on nine-species gene orthologs. Colors represent species. Panel titles show cluster identifiers and number of contained genes. **D** Developmental gene expression patterns in each of six clusters based on six vertebrate species gene orthologs. Panel titles show cluster identifiers and number of contained genes. **E** Relationship between the similarity of developmental gene expression patterns and phylogenetic distances. The expression similarity was calculated as the mean of Spearman correlation coefficients between mouse and non-mouse expression trajectories. **F** Chord graph indicating the relationship between clusters obtained using nine-species and six-species ortholog genes. Wider chords represent stronger connection between clusters.

The same analysis conducted using 2,038 development-related orthologs shared among six vertebrate species revealed a CL9.1-like developmental pattern represented by a single cluster (CL6.1) (Figure 3D). Genes in clusters CL9.1 and CL6.1 overlapped significantly and were enriched in similar GO terms (Figure 3F and Supplementary file 8: Table S5).

The second-largest cluster found in vertebrates (CL6.2) showed an opposite, ascending expression pattern conserved among species and contained genes enriched in signaling pathways, such as cGMP-PKG signaling pathway, oxytocin signaling pathway, and Renin secretion (Supplementary file 8: Table S5). Notably, this cluster overlapped with the chordate cluster CL9.4, where the ascending pattern was conserved among the seven species, excluding ciona and oyster (Figure 3C).

## Discussion

Our goal was to explore the general relationship between ontogeny and phylogeny based on the comparison of developmental gene expression trajectories among nine species separated by approximately 800 million years of evolution. Our results indicate that developmental patterns consistent with evolutionary models rooted in von Baer’s and Haeckel’s ideas match approximately half of all detected mRNA developmental expression profiles. Between the two historical concepts, the continuous developmental model consistent with Baer’s arguments was nearly three-fold more prominent compared to the progressive model compatible with the Biogenetic law. Still, a measurable fraction of genes displayed the developmental expression pattern matching Haeckel’s predictions, indicating the need for a further critical assessment of this historical concept. These results do not conflict with the hourglass model of development. Our previous work demonstrated the applicability of this model to all nine species’ development(Hu et al., 2017). While the hourglass model compares relative conservation of developmental stages, we focused on the alignment of expression differences among stages.

Expression profiles conserved along the entire species’ development included two main patterns: a gradual decrease conserved among all nine species and gradual increase conserved among six vertebrates. The conserved downward pattern coincides with the reported preferential expression of evolutionary older genes at earlier developmental stages(Gao et al., 2018). Consistently, genes forming this pattern tend to display ubiquitous expression among tissues and are involved in essential functions, such as RNA processing (Supplementary file 9: Figure S4). Further, a study examining embryonic gene expression reported the existence of such a pattern in isolated mouse tissues(Sarropoulos, Marin, Cardoso-Moreira, & Kaessmann, 2019), indicating that the descending expression reflects alterations within embryo tissues rather than changes in embryo composition. Overall, expression patterns conserved between species over the entire development involve 19-36% of orthologous development-related genes detected in our study, constituting the major alignment trend.

The second prominent type of developmental relationship among species, involving 5-15% of orthologous development-related genes, was generally consistent with predictions of Haeckel’s Biogenetic law. According to Haeckel’s views, the best developmental alignment should match stages reflecting the phylogenetic history of the species, rather than the entire developmental span. When aligning developmental profiles of eight species to the mouse, we indeed observed this phenomenon. First, for this gene group, the developmental expression profiles indeed aligned the best to the truncated mouse developmental series lacking late stages. Second, for each species, these alignments peaked at a particular shortened mouse developmental fragment, with species phylogenetically more distant from the mouse aligning best to the increasingly shorter mouse developmental sets. Intriguingly, while all species evolved their developmental programs after branching from the common ancestor, a mirror procedure, alignment to the fragmented developmental series of amphioxus and ciona representing more ancient body plans, revealed potential PM genes in only two of 16 alignments (Supplementary file 4: Figure S2). Further, there was no relationship between the length of the best-aligning developmental fragment and the phylogenetic distance between species.

Similar to the linearly-aligning genes, genes displaying this “Haeckelian” expression behavior overlapped significantly between species (Supplementary file 5: Table S3), indicating their potential functional conservation. These genes were indeed more conserved at the amino acid sequence level than the linearly-aligning genes and displayed enrichment in specialized functions, such as synaptic signaling and secretion.

Our work highlights the usefulness of quantitative endophenotypes as an essential complement to morphology-based developmental studies. It further suggests partial applicability of Haeckel’s ideas to a fraction of developmental expression differences involving genes associated with neural functions.

## Methods

### RNA-seq data processing

All data in this analysis were downloaded from Hu et al.(Hu et al., 2017). We adopted the same methods to quantify gene expression. In brief, we got the RNA-seq data of *Crassostrea gigas* from GEO database (accession number SRP014559)(Zhang et al., 2012), and the rest of species from DDBJ (accession number DRA003460) (Supplementary file 1: Table S1). To quantify gene expression, we mapped RNA-seq reads to the corresponding genome (Supplementary file 1: Table S1) using Tophat (v2), allowing up to three mismatches and indels, except for ciona (Ci). In the case of ciona, we mapped reads to the genome with up to five mismatches and indels as the ciona RNA-seq data and the genome data have slightly lower quality compared to the rest.

To define the expressed genes, we required the maximal expression across all development stages exceeding 1 FPKM. For each stage, we calculated expression as the mean of expression of the replicated samples. More than 70% of coding genes annotated for a given species were reliably detected, except for amphioxus (41%) (Supplementary file 10: Table S6).

### Identification of development-related genes

We defined developmental alterations of gene expression levels using polynomial regression models following the method described in Somel et al.(Somel et al., 2009). For each gene, we chose the best regression model with the developmental stage (by rank) as predictor and expression level as a response with Benjamini-Hochberg corrected *p* < 0.05. The genes that fit a significant regression model were termed as development-related genes (Figure 1D -Source data 1).

### Identification of stage-associated genes

We defined genes preferentially expressed at a particular developmental stage as stage-associated genes in each species following the method described in(Hu et al., 2017; J. J. Li et al., 2014). For each species, we required FPKM (fragments per kilobase of exon model per million reads mapped) > 2 at a particular stage and Z-score (normalized FPKM across samples) representing the difference between this stage and the rest of stages > 1.5. On average, 65%-90% of genes expressed in development were identified as stage-associated genes (Supplementary file 2: Table S2).

### Alignment of mouse developmental stage sets to developmental profiles of other species

To assess the transcriptome similarity in the course of developmental stages between mouse (Mm) and other species, we used stage-associated and 1:1 orthologous genes to align the developmental stages between mouse and other species followed by the method reported in(Hu et al., 2017; J. J. Li et al., 2014). The detailed description can be found in the supplementary materials of(Hu et al., 2017; J. J. Li et al., 2014). In brief, we calculated the pairing score using the hypergeometric test to evaluate the ratio of orthologs overlapping within the stage-associated genes pairwise. The pairing score can be written as:

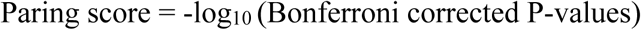

The paring score was used to quantify the significance of the overlap on each pairwise stage comparison between the mouse and another species. From the paring score, we identified the relationship between mouse and other species (Figure 1E). To check the stability of this relationship, we repeated the comparison 500 times with randomly assigned stage-associated genes for each stage of each species using the same procedure.

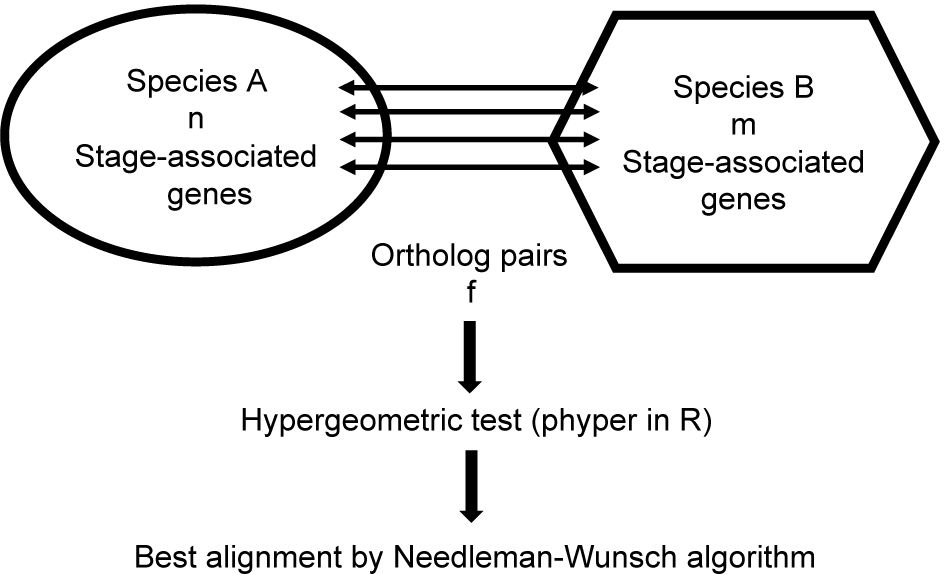

According to the above scheme, we further assigned the corresponding stage alignment between mouse and other species using the Needleman-Wunsch algorithm with the gap penalty equal to one. To estimate the stability of alignment based on the mean of the replicates, we randomly chose one individual per stage, aligned two species 500 times, and calculated the frequency for each pair of alignment. The resulting frequency was presented by the thickness of the line in Figure 3A.

### Identification of CM/PM genes

To identify genes with expression trajectories compatible with the continuous model (CM) or progressive model (PM) predictions, we considered development-related genes as mentioned in “*Identification of development-related genes*” with existing orthologs between mouse and any other species. We then linearly aligned the complete set of developmental stages in a given species (oyster, amphioxus, ciona, zebrafish, frogs, turtle, and chicken) to the increasingly truncated sets of mouse developmental stages, i.e., from first two to all 17 stages (Figure 2A–figure supplement 5). To this end, we interpolated 20 time points uniformly distributed across the whole ontogeny of non-mouse species using cubic smoothing spline (with three degrees of freedom for amphioxus data and six degrees of freedom for other species) or polynomial regression curves (up to the fourth degree). The two methods for interpolating expression values across development were used to ensure the robustness of our CM or PM predictions. Of note, during cubic smoothing spline, the degrees of freedom chosen for amphioxus were fewer than other species, due to the relatively fewer sampled developmental stages of this species; whereas during polynomial regression, we chose up to four degrees to take advantage of the near-fitness of polynomial regression in most time series data.

We then aligned the resulting expression trajectories to the gradually increasing sets of mouse developmental stages, where the expression trajectory was interpolated using the same approach (cubic smoothing spline with six degrees of freedom or polynomial regression with up to the fourth degree). The alignments with the Pearson correlation coefficient (PCC) greater than 0.7 and correlation test *p*-value less than 0.05 were considered as valid. We further selected the alignment with the maximum PCC among all valid alignments as the ultimate one. Genes failing to pass the two alignment criteria were considered as non-aligned, regardless of the complete or partial linear alignment paradigms.

Based on the above classification procedure, we summarized for each species, the number of genes aligning to each set of developmental stages of the mouse. Furthermore, we repeated the classification process by shuffling the orthologous relationships between non-mouse species and the mouse (Figure 2B–figure supplement 6). Specifically, we expected that by shuffling, each developmental stage of the mouse would be equally represented in other species, allowing us to calculate the random expectation of alignment occurrences for each stage. Thus, the final number of genes aligning to each set of developmental stages of the mouse was obtained after subtracting these background contaminations. Genes adhering to the continuous model (CM) were defined as those best aligned between mouse and other species over the complete developmental intervals. By contrast, genes following the progressive model (PM) were defined by the following criteria: i) aligned best to a truncated set of mouse developmental stages; ii) displayed a significantly greater number of aligning genes compared to the background distributions, determined using a BH-corrected *p*-value of less than 0.05 as a cutoff; and iii) an increase in the number of genes aligning to particular sets of mouse stages compared to its adjacent sets (Figure 2A, Figure 2E – source data 1 and Figure 2E – source data 2).

To further check the robustness of the alignment results, we restricted our analysis to evolutionarily old genes. To do so, we repeated the alignment procedure using a subset of the development-related genes with inferred Earliest Ortholog Level (EOL) < level 10. This conservation level corresponds to genes that appeared and became fixed within the genomes before the separation of Chordata(Litman & Stein, 2019).

### Overlap of CM/PM genes between species

To calculate the significance of the overlap of CM or PM genes between species, we sampled the same number of CM or PM genes in each species from all development-related genes 1,000 times and recalculated the number of overlapping genes to obtain the empirical distribution. The overlap significance *p*-value was calculated as the proportion of the values, which were as larger or greater than the actual overlapping gene number. Given all pairs of species involved, a Bonferroni-corrected *p*-value of less than 0.05 was used as the cutoff of the overlap significance (Supplementary file 5: Table S3).

### Amino acid conservation of CM/PM gene

We compared the evolutionary conservation between proteins encoded by CM and PM genes using the Ka/Ks metric(Yang & Nielsen, 2000). Since in this study, we are not studying Ka/Ks variations across the evolutionary branches – we focused on the comparisons of PM versus CM genes within each individual species, we conducted this analysis mainly in a pairwise manner between a given species and mouse. Firstly, we extracted the coding sequences of each gene based on the corresponding annotation information and then translated the sequences into amino acid sequences using the function “translate” implemented in the Bioconductor package “Biotrings". The longest protein sequence was selected as the gene protein sequence. Next, we performed protein sequence alignments between non-mouse species and the mouse using the function “pairwiseAlignment” in “Biostrings", with the scoring matrix set as BLOSUM50. The resulting protein alignment was translated to the nucleotide level and used as the input for PAML(Yang, 1997) for a Ka/Ks analysis output. This procedure was conducted in a pairwise manner for the mouse and a given non-mouse species to match the corresponding definitions of PM and CM genes carried out in a pairwise manner (Figure 2G–figure supplement 8). To test the robustness of our Ka/Ks calculations in the context of multiple species, we performed multiple sequence alignment using Clustal Omega(Sievers et al., 2011) for all protein sequences mentioned above convert to nucleotide levels. We next utilized the CODEML application in PAML to calculate the Ka/Ks for given cross-species ortholog. The significance of the difference between PM and CM genes in each species was assessed using one-sided Wilcoxon rank-sum test (Figure 2G–figure supplement 9).

### Gene expression interpretation to mouse developmental stage

To compare the similarity of the expression profiles across developmental stages of nine species, we used the predicted developmental stage alignment presented in Figure 3A to create a unified alignment of eight species to the mouse developmental curves. To do so, we mapped 33 stages cumulatively interpolated from eight species to the full mouse developmental curve fitted using cubic smoothing spline with ten degrees of freedom. We then compared gene expression curves among nine species based on z-transformed expression of each gene interpolated at these 33 stage points.

### Clustering of gene expression profiles in six vertebrate and nine chordate species

To investigate the expression pattern diversity in nine or six species, we used hierarchical clustering (hclust function in R) of z-transformed gene expression trajectories aligned among species with (1 - rho) as the distance measure, where rho is the Spearman correlation coefficient. We chose k equal six, as optimal, based on visual inspection of clusters obtained using different k values.

### Functional annotation of developmental expression patterns

To check the functions of genes in each cluster, we applied Gene Ontology (GO) and Kyoto Encyclopedia of Genes and Genomes (KEGG)(Kanehisa & Goto, 2000) enrichment tests. GO annotation of mouse genes was downloaded from Bioconductor package ‘org.Mm.eg.db’. Mouse pathway annotation was downloaded from http://rest.kegg.jp/list/pathway/ and http://rest.kegg.jp/link/genes/.

For GO enrichment test, the “elim” algorithm of topGO(Alexa & Rahnenfuhrer, 2016) was chosen to eliminate the hierarchical dependency of the GO terms. Fisher’s exact test was applied for each GO term. The background set consisted of all development-related genes orthologous among six vertebrate species (2,038 genes) and nine chordate species (244 genes), respectively. For the test of vertebrate gene clusters, Benjamini-Hochberg correction on “molecular function”, “biological process”, and “cellular component” was applied independently. GO terms with BH corrected *p* < 0.05 were reported. We applied no multiple test correction to chordate’ gene cluster enrichment analysis due to low statistical power of the dataset and used a more stringent nominal *p*-value cutoff of *p* < 0.001.

For the pathway enrichment test, the reference genes were the same as for GO-based analysis. We used hypergeometric test (phyper in R) to assess the enrichment in each KEGG pathway. Bonferroni correction was applied for genes in vertebrate clusters. Pathways with corrected *p* < 0.05 were reported as significantly enriched. No correction was applied to genes from chordate clusters due to low statistical power, and the nominal pathway enrichment *p*-value was set to *p* < 0.05. This relaxed cutoff was used to assess the potential overlap between enriched functions found in vertebrate and chordate clusters (Supplementary file 8: Table S5).

### Species tree construction

The species tree was generated under the web https://phylot.biobyte.de/ with NCBI taxonomy IDs (10090, 9031, 13735, 8364, 7955, 7719, 8355, 7739, 29159). The species separation time to the mouse was obtained from http://www.timetree.org/(Kumar, Stecher, Suleski, & Hedges, 2017).

### Statistical analysis and software

Analyses were conducted in the R environment (http://www.r-project.org/). To minimize the type I error rate, we used multiple test correction for *p*-value calculations, unless specifically indicated otherwise. The statistically significant level used for each was specified in the main text. Additionally, we used Perl, Python, and R packages, including ‘topGO’, ‘reshape’, ‘RColorBrewer’, ‘ggplot2’,’ Biotrings’, as well as shell scripts, in the analyses. Pathway visualization was conducted under https://pathview.uncc.edu/ for the spliceosome pathway.

## Acknowledgments

We thank Michael Lachmann for inspiring this study, Zhisong He, Qian Li and Qianhui Yu for helpful discussions, and Pavel Mazin for critical comments.

## Author contribution

S.G, HY.H and C.X designed and executed the bioinformatics analysis. S.G and PK wrote the manuscript. N.I and P.K revised the manuscript. PK supervised the project.

---

**Figure 1A–figure supplement 1.**
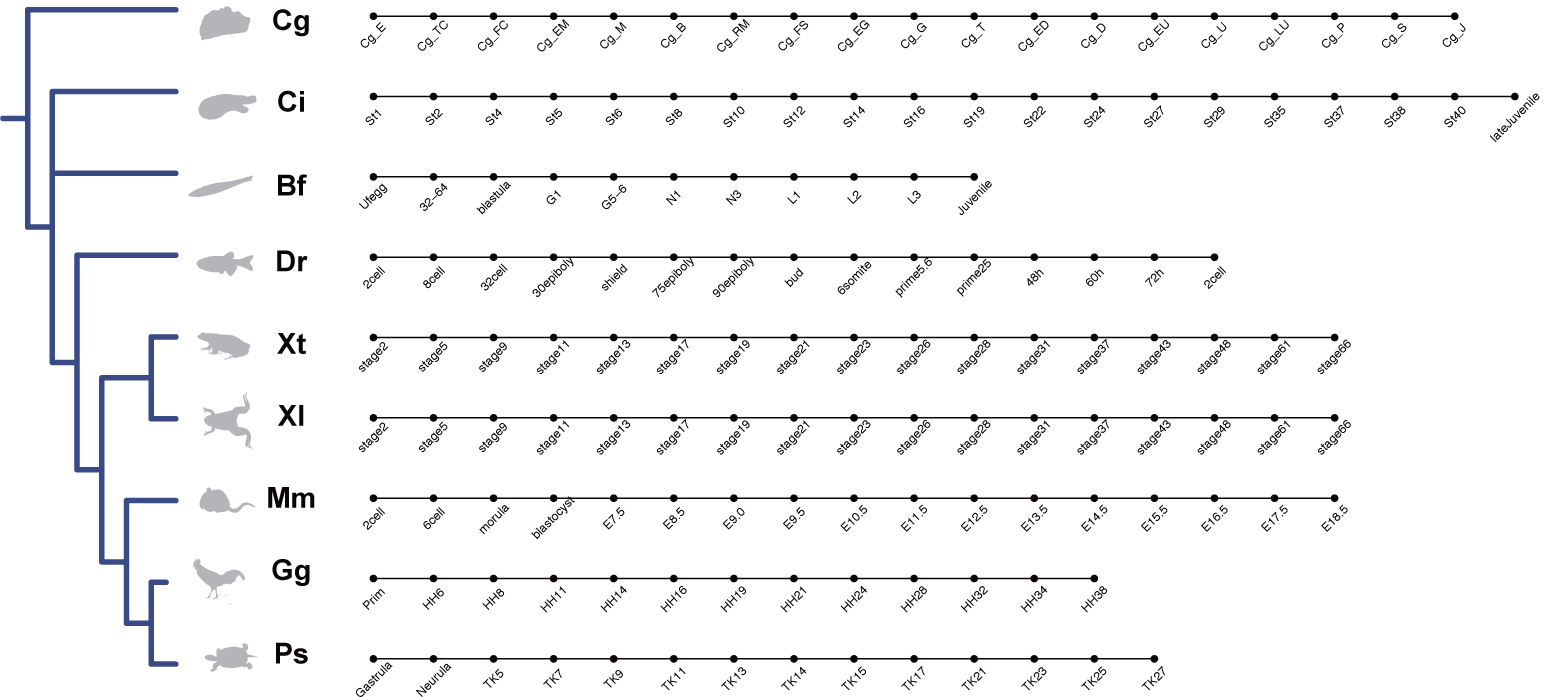
Illustration of sampled developmental stages for each species correspondent to Figure 1A. The dendrogram shows the phylogenetic relationship among species, indicated by silhouette figures and two-letter abbreviated species’ classification names. Dots indicate sampled embryonic stages. Detailed sampled stage information is listed in Supplementary file 1: Table S1.

**Figure 1C–figure supplement 2.**
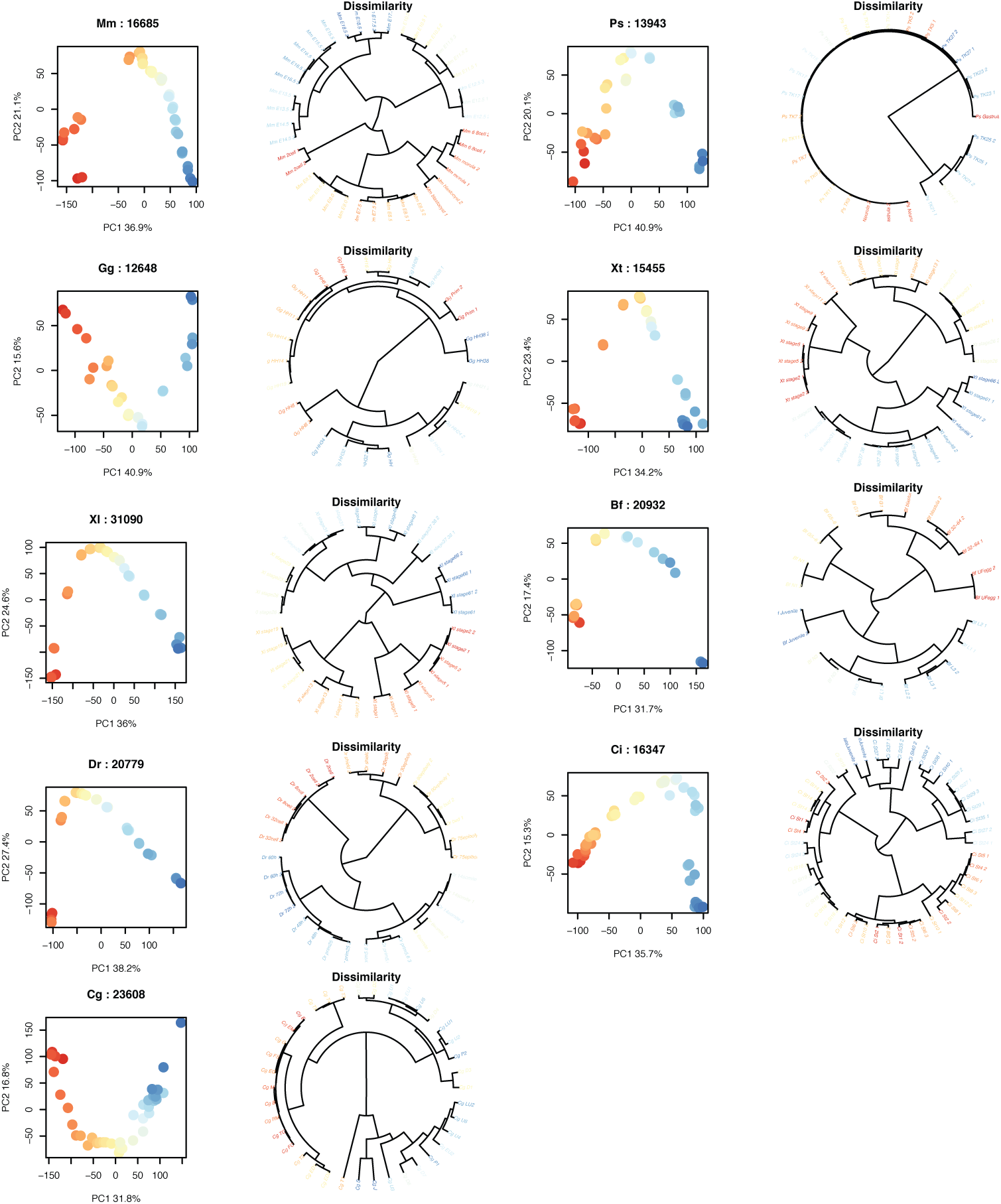
PCA plot and dendrogram based on expression variation within each species. Dots represent samples, color represents developmental stages (red – early, blue – late). Dendrograms were constructed using 1-Spearman correlation coefficient dissimilarity matrix. Panel titles show abbreviated species’ identifiers and the number of expressed genes with FPKM > 1 in at least one developmental stage.

**Figure 1E–figure supplement 3.**
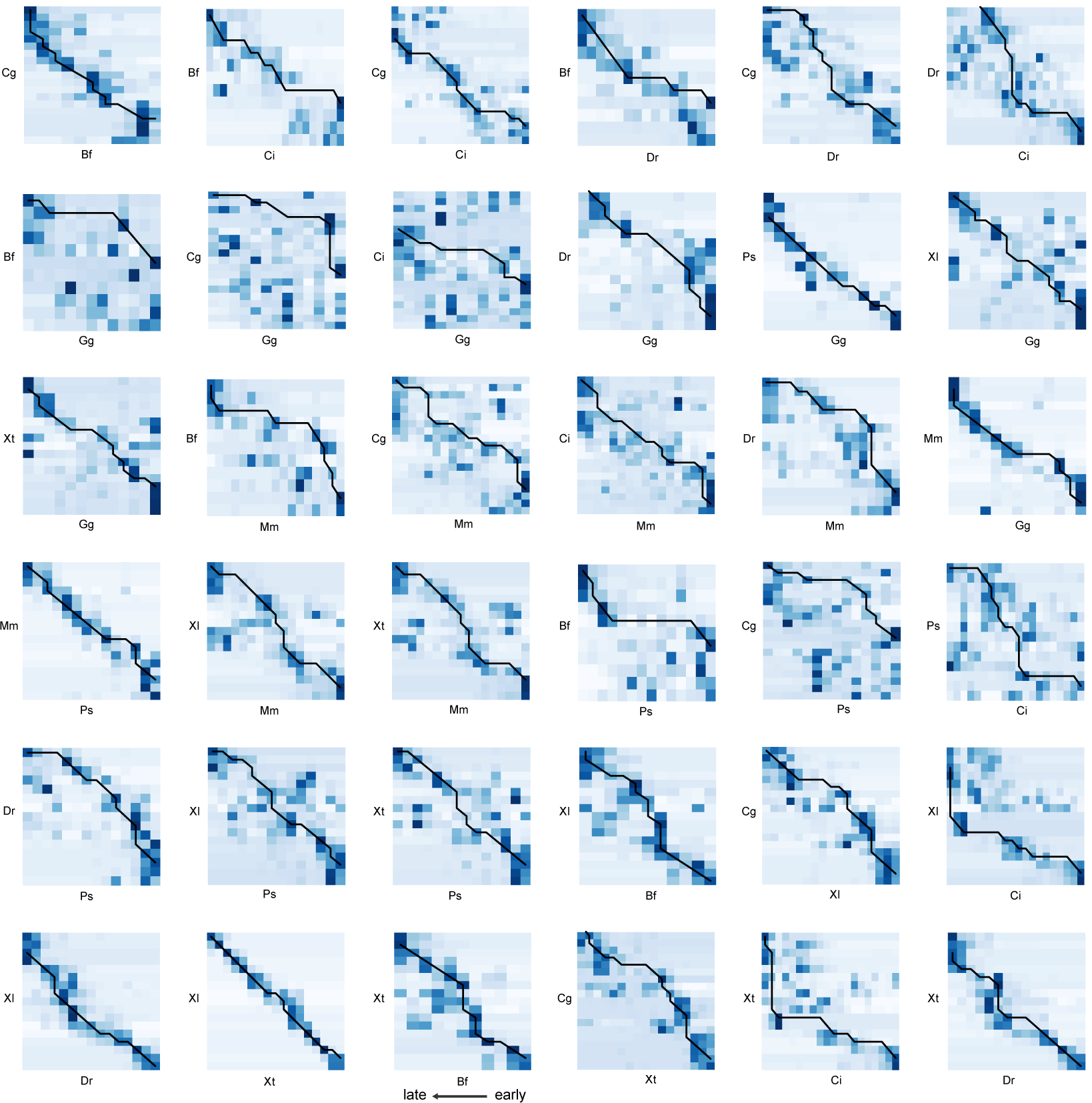
Heatmap of the pairing scores reflecting the overlap of stage-associate genes correspondent to Figure 1E. Darker shade of blue represents greater overlap of stage-associate genes. Black line represents the optimal alignment path calculated using the Needleman-Wunsch algorithm.

**Figure 1E – figure supplement 4.**
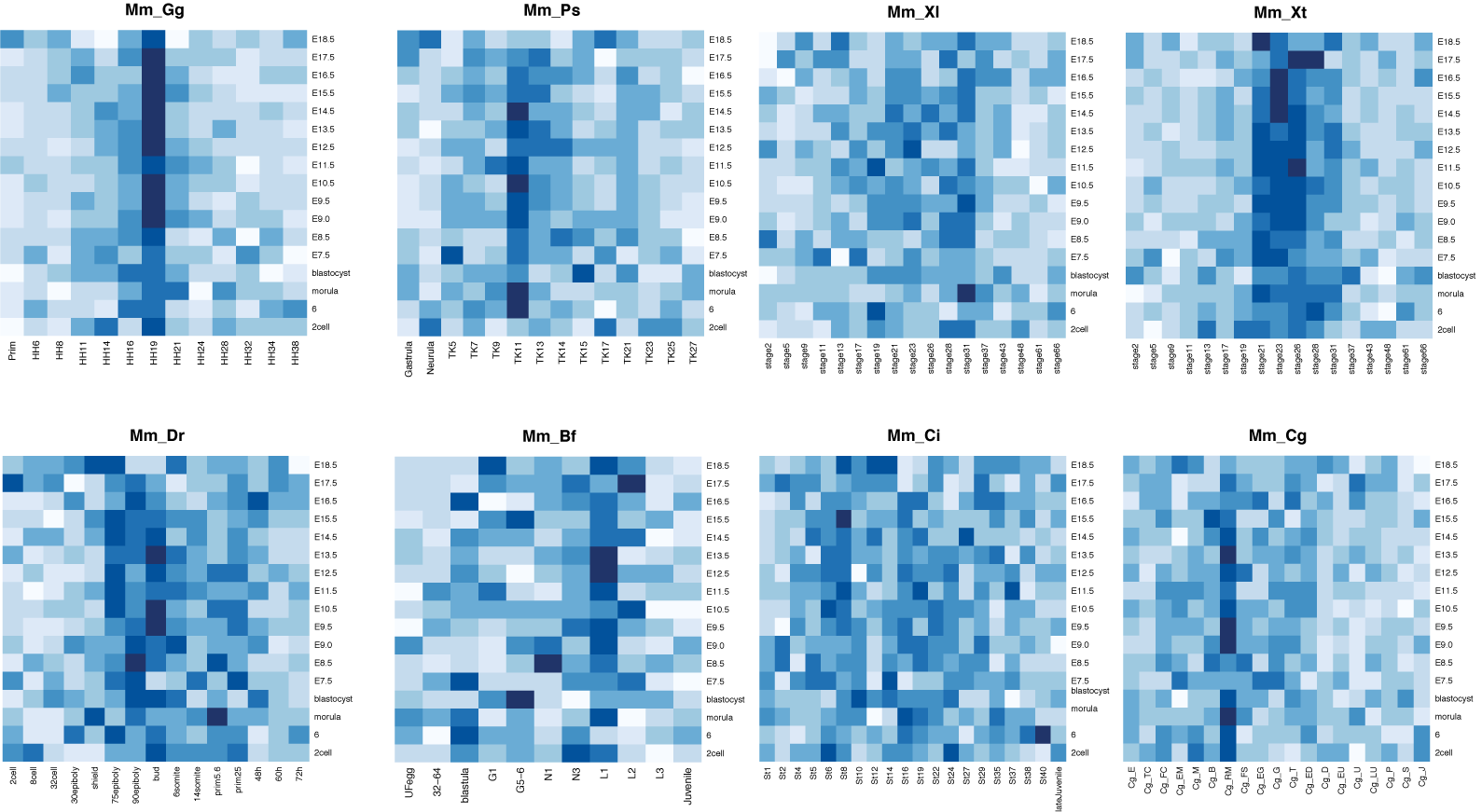
Heatmap of the average pairing scores calculated randomly assigned stage-associated genes correspondent to Figure 1E. The heatmap shows pairing scores based on the average of 500 iterations of random assignments of stage-specific gene labels. Panel titles show abbreviated species’ identifiers. Darker shades of blue represent greater overlap between stages.

**Figure 2A – figure supplement 5.**
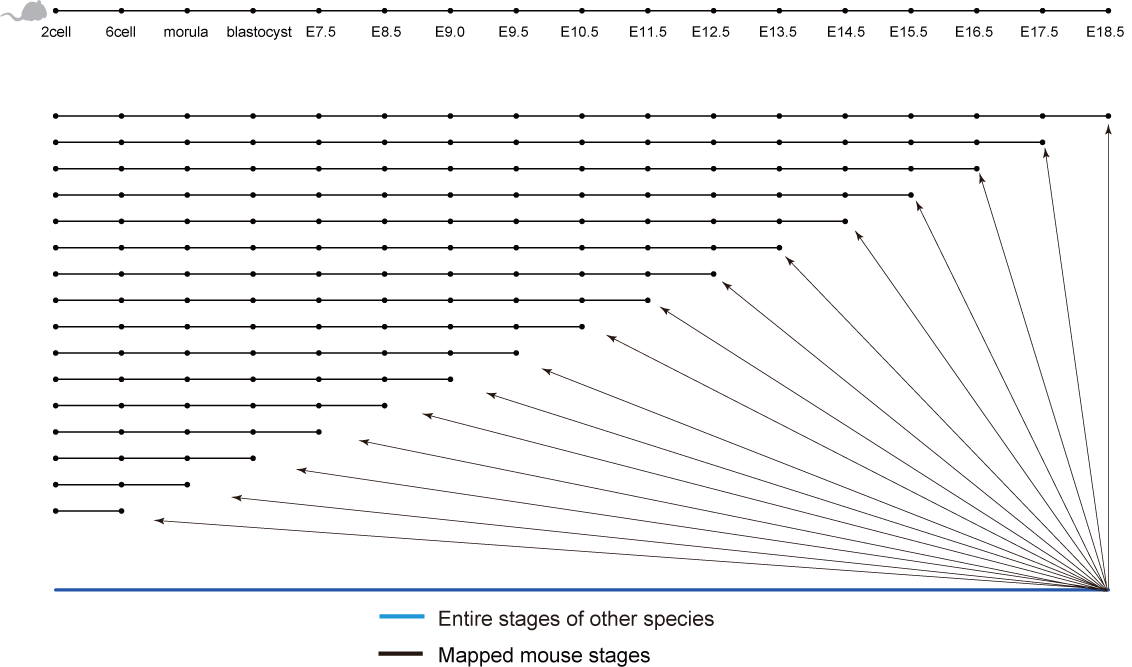
Schematic representation of the alignment procedure used to test the predictions of progressive and continuous models correspondent to Figure 2A. The scheme depicts an example of all possible alignments between the complete developmental series of a species (any of investigated species except mouse; blue line) and mouse developmental sets (black lines). The mouse developmental sets included the entire developmental series consisting of 17 sampled stages and developmental series progressively shortened by truncation of the last stage, down to the first two developmental stages. For each gene showing development-dependent expression in the two compared species, we then aligned the developmental profile of a species to the entire mouse developmental course and the increasingly truncated sets of mouse developmental stages. The continuous model (CM) predicts the best alignment of the amphioxus development to the complete mouse developmental course. The progressive model (PM) predicts the best alignment to the truncated developmental course. We used all possible truncated mouse developmental series, from 16 to two stages, to identify the best alignment to the mouse developmental series in an unbiased manner.

**Figure 2B – figure supplement 6.**
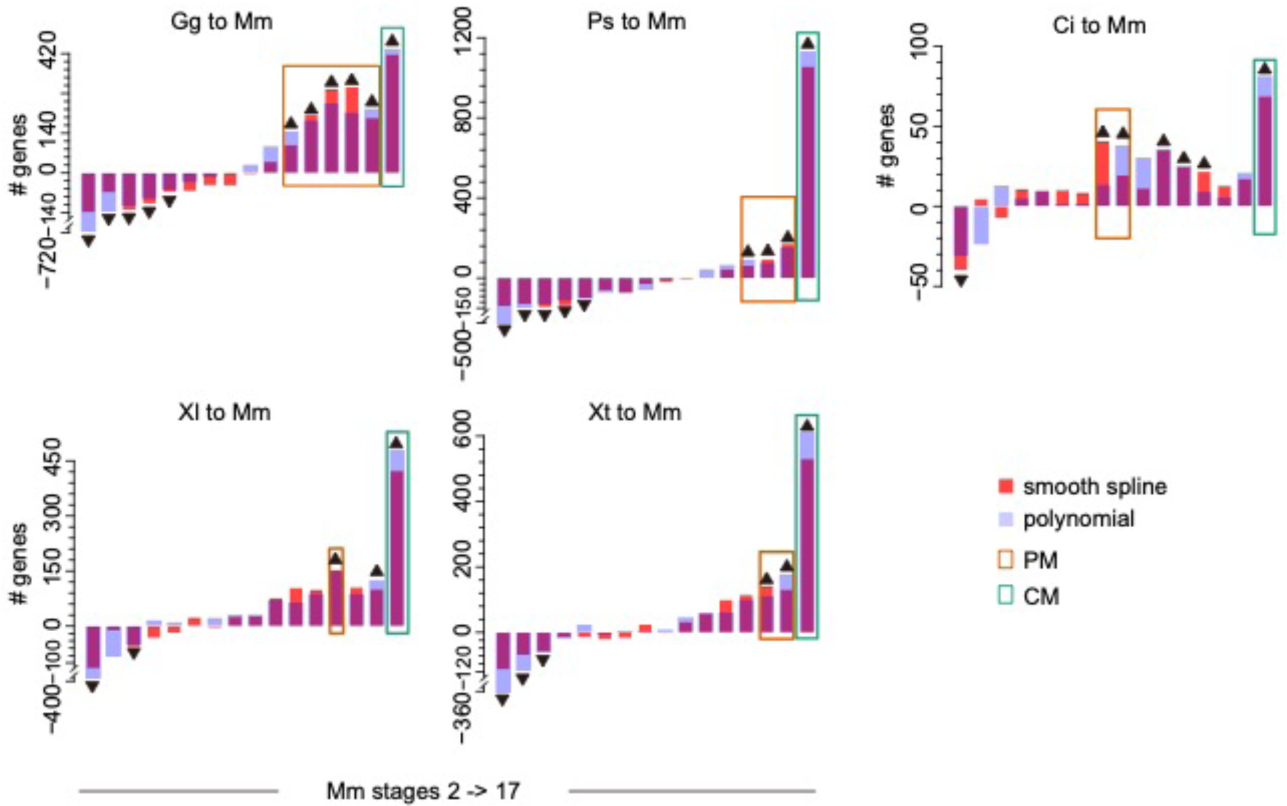
Numbers of genes showing the best expression trajectory’s alignment between the complete developmental course of five species and mouse developmental sets of different lengths: from two to 17 stages. Panel titles show abbreviated species’ identifiers. Colors indicate two methods used for developmental expression trajectory calculation: cubic smooth spline (orange) and polynomial regression (gray). Colored rectangles mark stages containing alignments fitting CM (green) or PM (orange) predictions. Black triangles indicate significantly greater (up) or lesser (down) number of genes than that expected by chance aligning to an indicated mouse developmental interval (permutation test, BH-corrected *p* < 0.05). The background distribution of gene numbers for subtraction was estimated by shuffling the orthologous relationships of mouse and a given species.

**Figure 2C – figure supplement 7.**
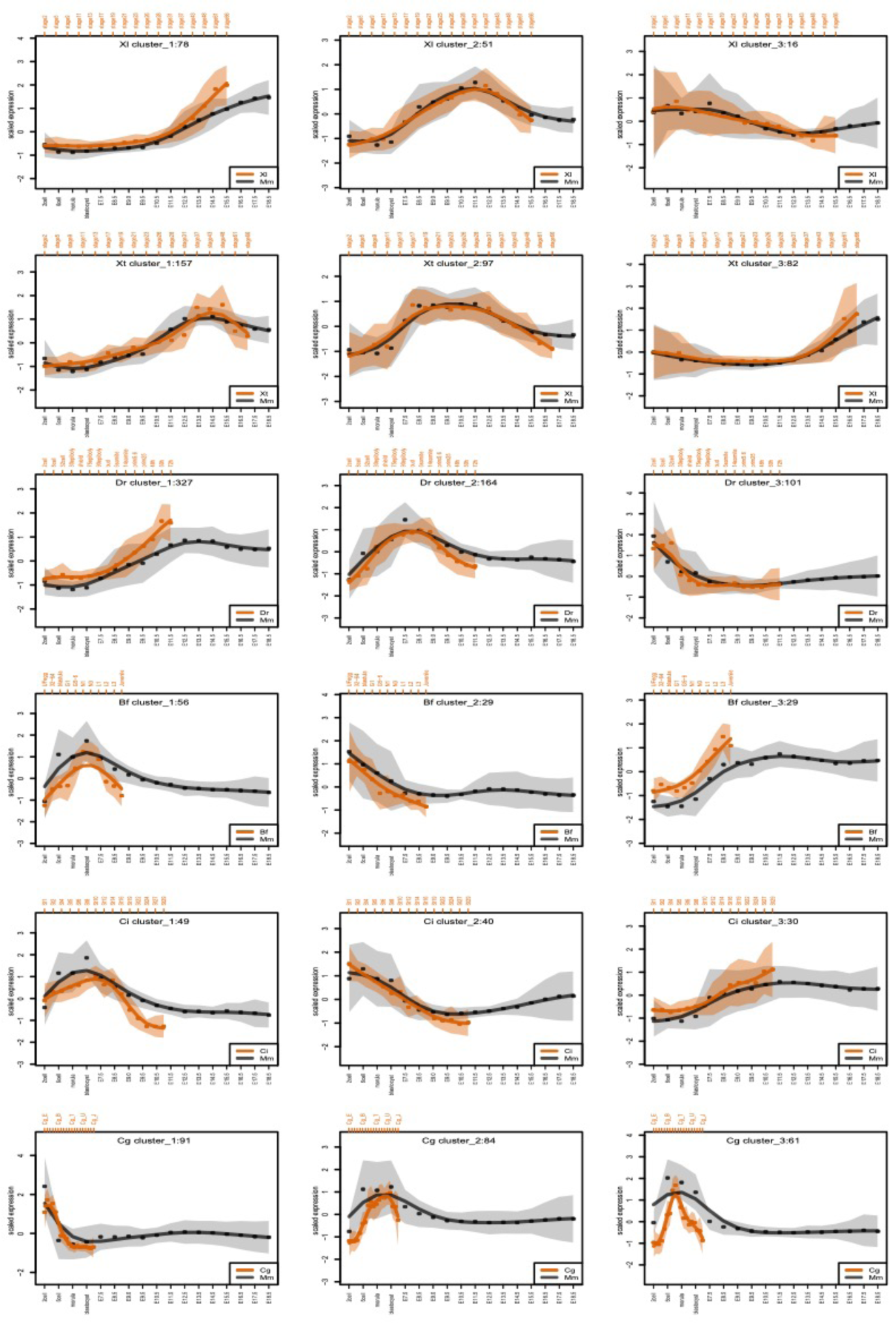
Examples of PM gene expression patterns correspondent to Figure 2C. Dots show the cluster-level standardized expression levels at each developmental stage in mouse (black) and the other species (orange). The curves represent average expression profiles and the shaded regions represent the standard deviation of curve estimates.

**Figure 2G – figure supplement 8.**
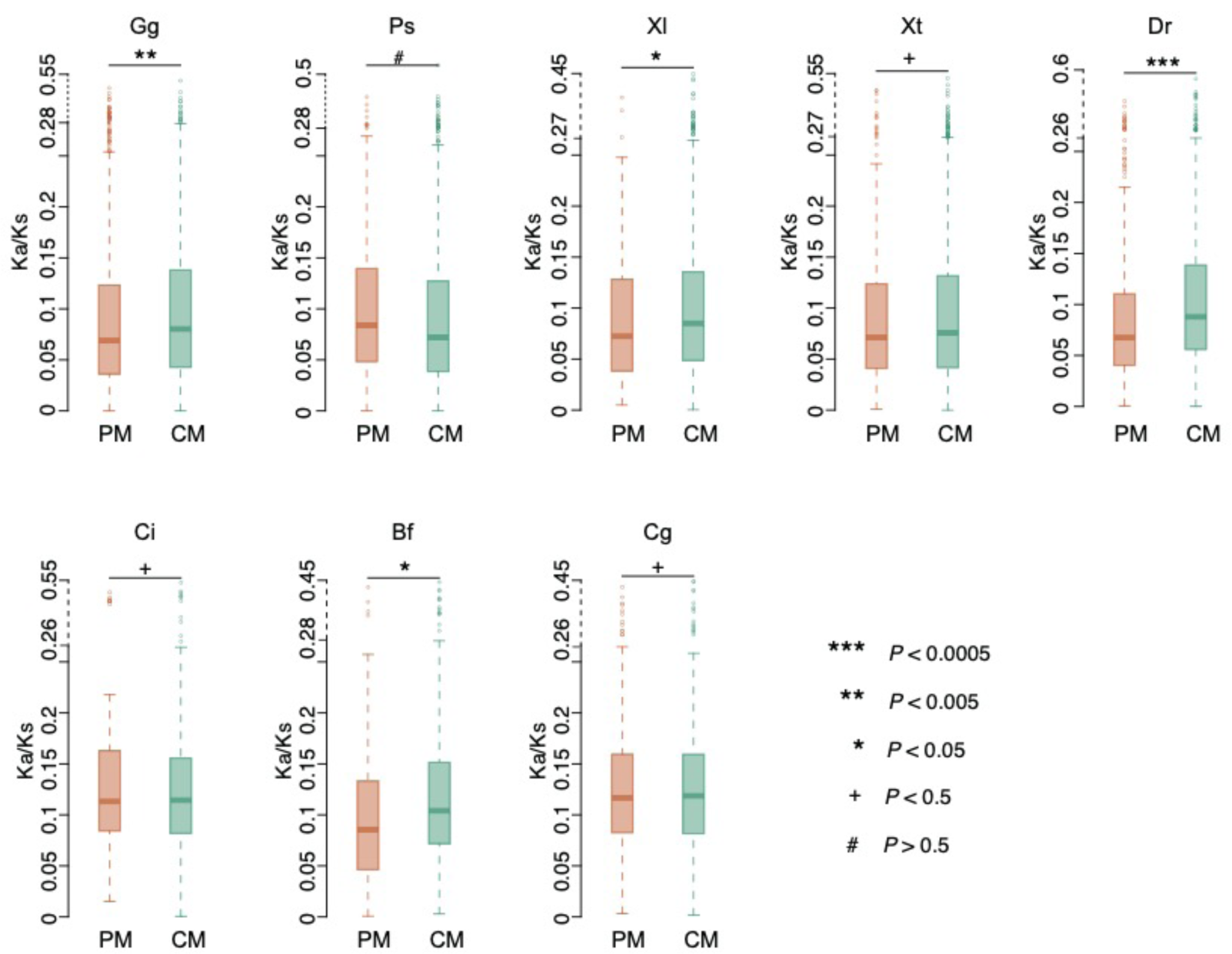
The distributions of Ka/Ks values of PM and CM genes in each non-mouse species correspondent to Figure 2G. Significance of the difference between the two distribution was assessed using one-sided Wilcoxon rank-sum test. ***, *p* < 0.0005, **, *p* < 0.005, *, *p* < 0.05, +, *p* < 0.5, #, *p* > 0.5.

**Figure 2G – figure supplement 9.**
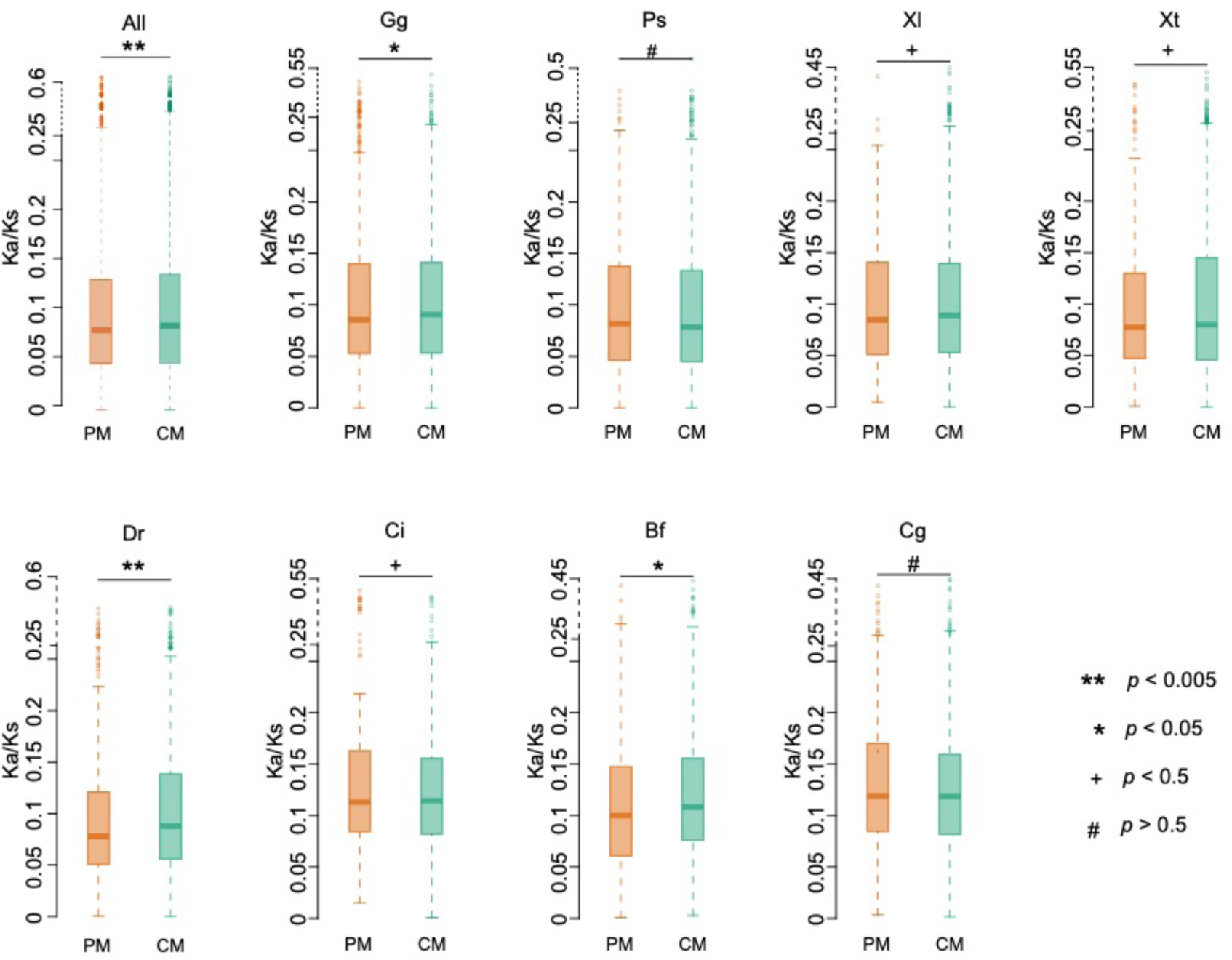
The distributions of Ka/Ks values of PM and CM genes in all species, and in each of the non-mouse species correspondent to Figure 2G. Significance of the difference between the two distributions was assessed using one-sided Wilcoxon rank-sum test. ***, *p* < 0.0005, **, *p* < 0.005, *, *p* < 0.05, +, *p* < 0.5, #, *p* > 0.5. In contrast to results displayed in Figure 2G and Figure 2G – figure supplement **8**, the Ka/Ks ratios here were estimated using multiple species alignment.

**Figure 3E – figure supplement 10.**
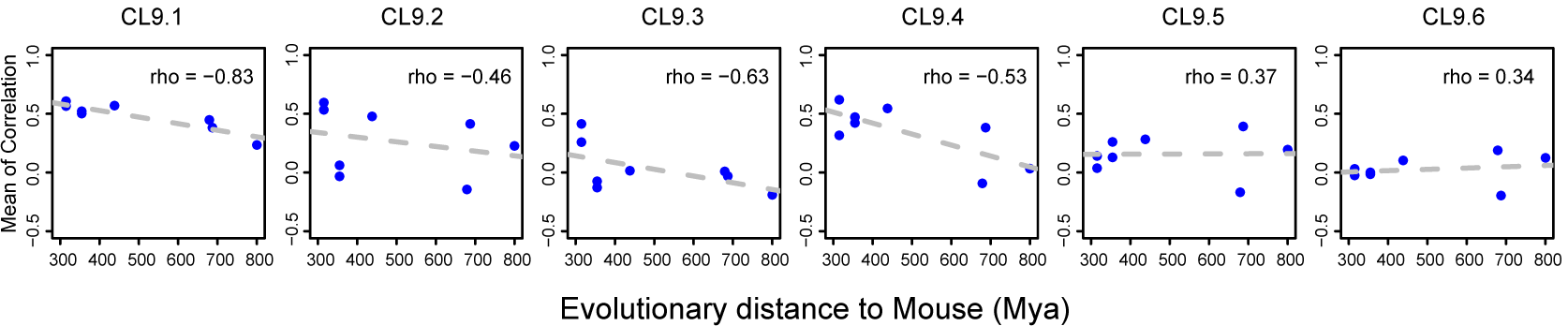
Relationship between the similarity of developmental gene expression patterns and phylogenetic distances correspondent to Figure 3E. The relationship is shown for each of six clusters of development-related genes orthologous among nine species. The expression similarity was calculated as the mean of Spearman correlation coefficients between mouse and non-mouse expression trajectories. The x-axis shows the phylogenetic distance to the mouse.

**Supplementary file 3: Figure S1.**
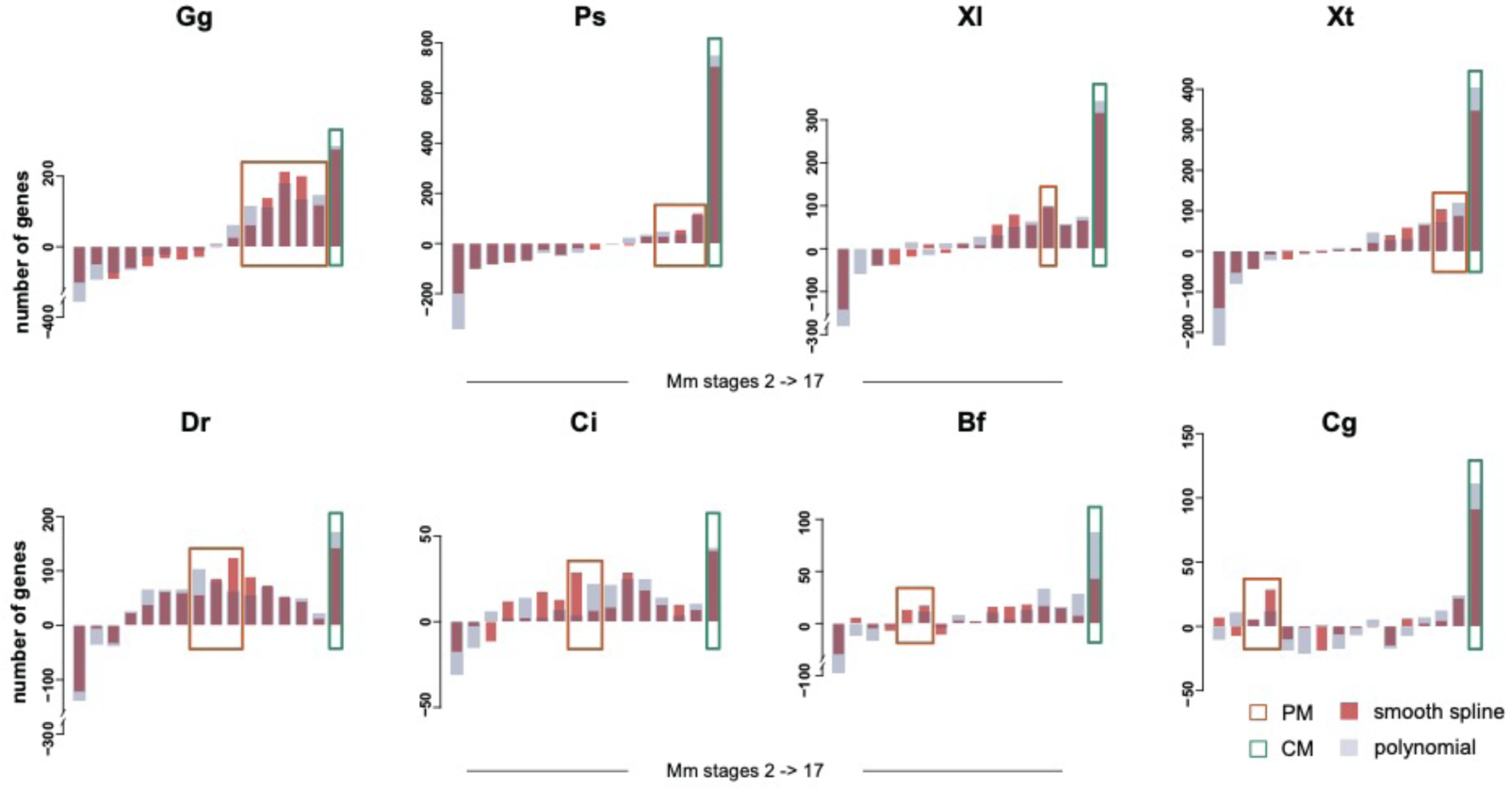
Numbers of genes showing the best expression trajectory’s alignment between the complete developmental course of eight species and mouse developmental sets of different lengths: from two to 17 stages. Panel titles show abbreviated species’ identifiers. Bar colors indicate two methods used for developmental expression trajectory calculation: cubic smooth spline (orange) and polynomial regression (purple). Colored rectangles mark stages containing alignments fitting CM (green) or PM (orange) predictions as in Figure 2B. The analysis was restricted to a subset of evolutionarily old genes inferred to exist back in *chordata* ancestor or earlier.

**Supplementary file 4: Figure S2.**
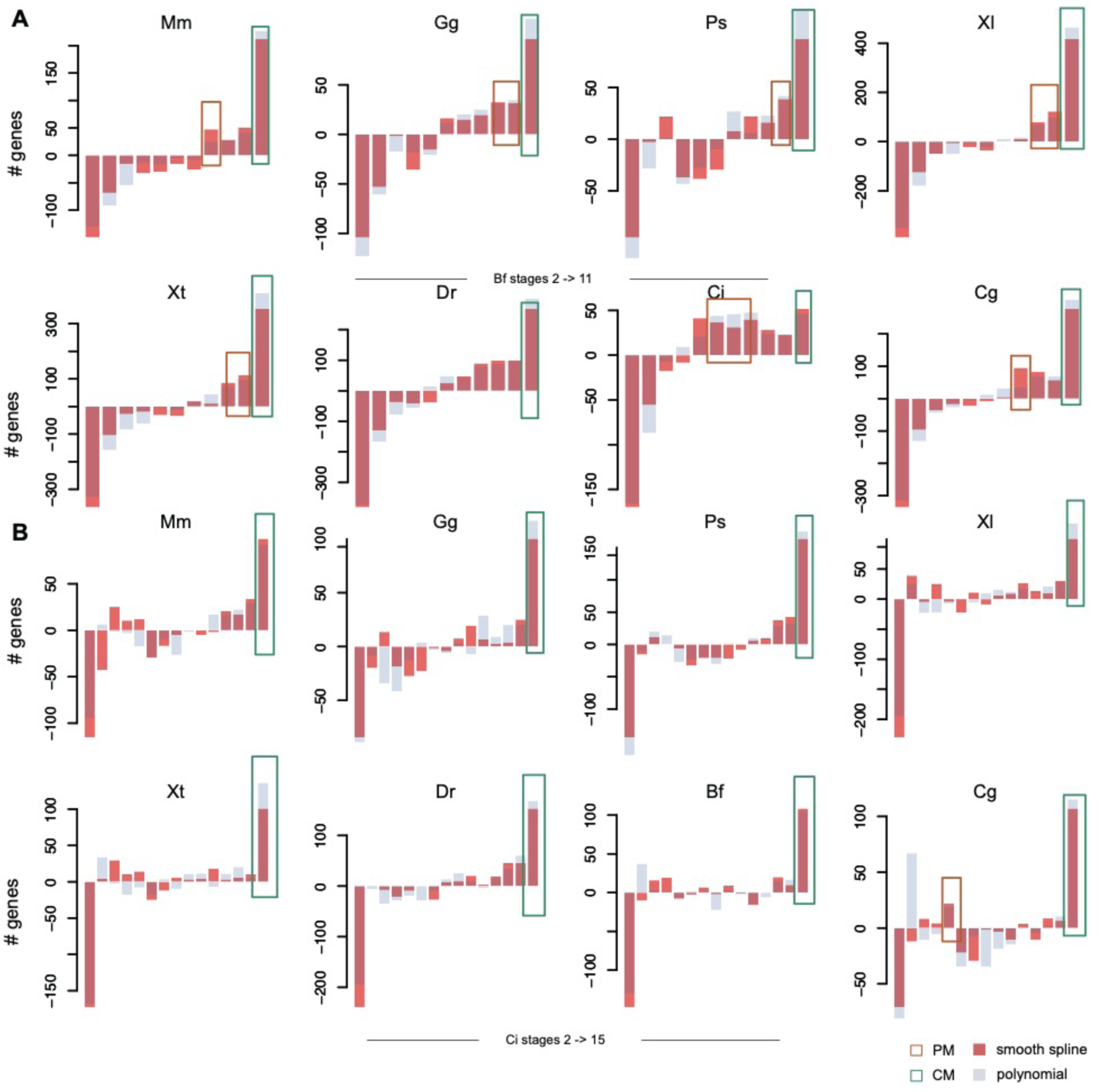
Numbers of genes showing the best expression trajectory’s alignment between the complete developmental course of eight species and amphioxus (panel A) and ciona (panel B) developmental sets of different lengths. Panel titles show abbreviated species’ identifiers. Bar colors indicate two methods used for developmental expression trajectory calculation: cubic smooth spline (orange) and polynomial regression (purple). Orange and green colored rectangles mark stages corresponding to alignments with significant excess of genes following PM and CM predictions respectively, as in Figure 2B.

**Supplementary file 6: Figure S3.**
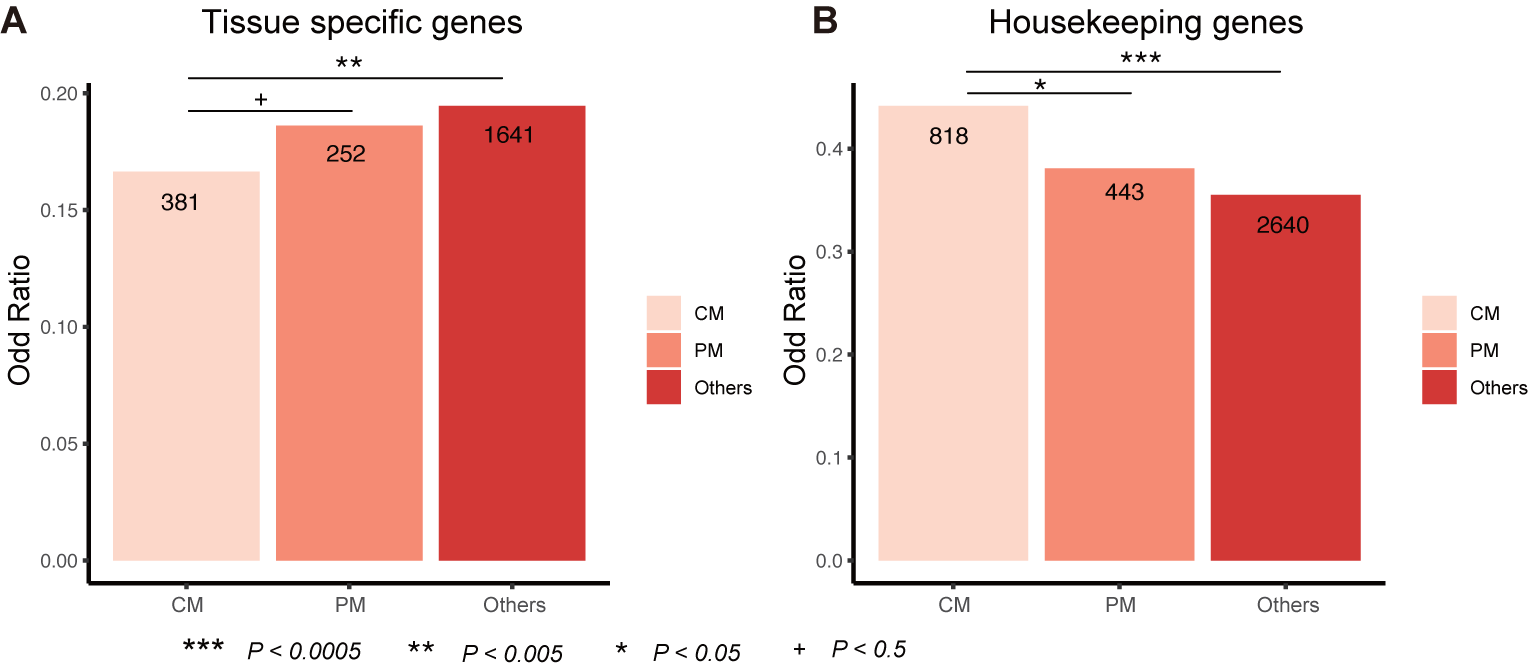
Tissue specificity of CM and PM genes. **A** Bars represent the ratio of tissue-specific and non-tissue specific genes for three gene groups. Numbers within bars show the number of tissue-specific genes in each group. **B** Bars represent the ratio of housekeeping and non-housekeeping genes. Numbers within bars show the number of housekeeping genes in each group. The significance of the differences between gene groups was assessed using one-sided Fisher’s exact test. ***, *p* < 0.0005, **, *p* < 0.005, *, *p* < 0.05, +, *p* < 0.5.

**Supplementary file 9: Figure S4.**
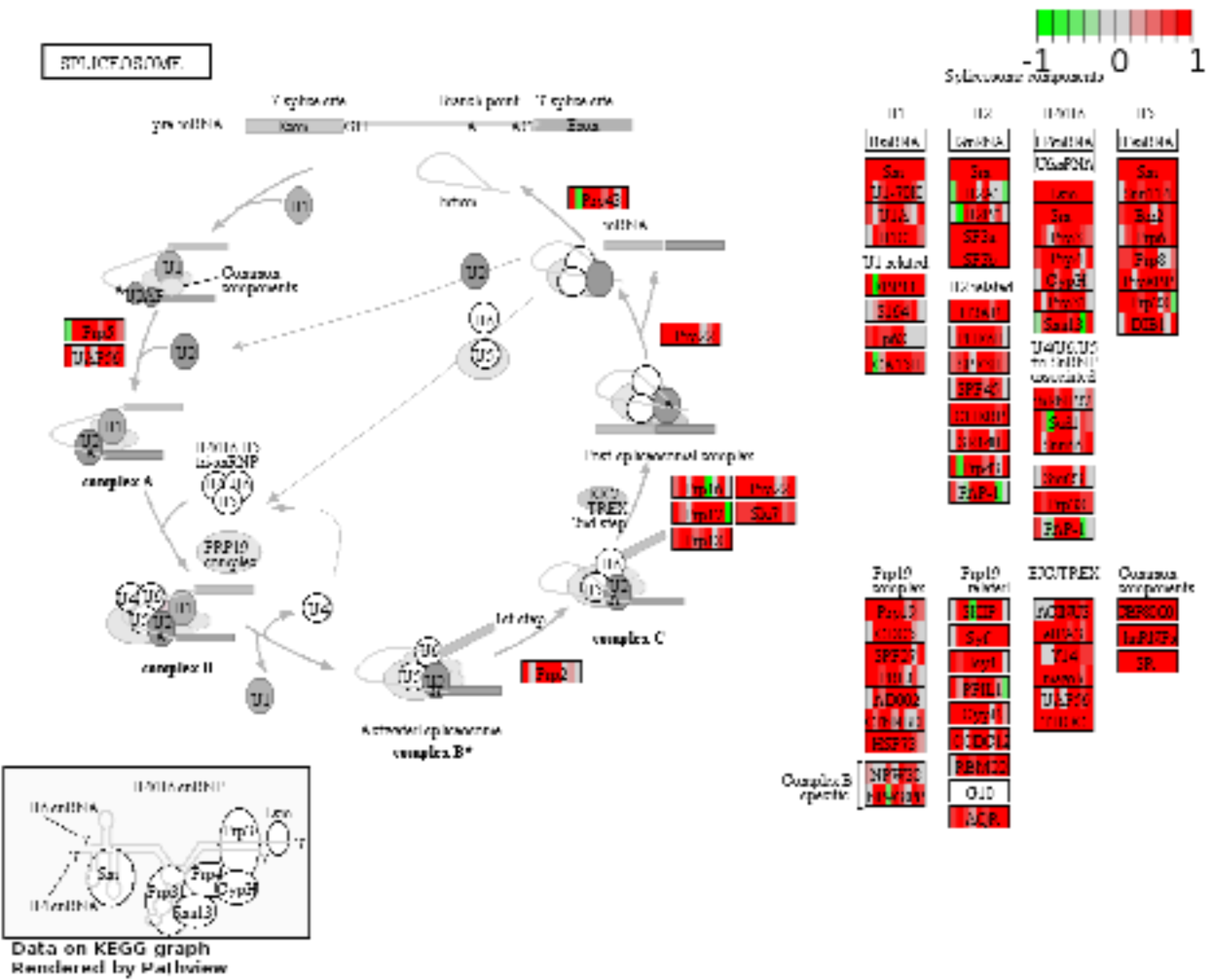
Visualization of spliceosome pathway showing developmental expression profile correlation among nine species. Scheme shows mouse orthologs of spliceosome pathway genes present in nine species. Colors indicate the coefficients of developmental expression profile correlations estimated in all pairwise comparisons between mouse and the other eight species.

## Additional files

**Figure 1D – source data1. Developmental related gene list of each species correspondent to Figure 1D**.

**Figure 2E – source data1**. The list of continuous model genes for each species

**Figure 2E – source data2**. The list of progressive model genes for each species

**Figure 2F – source data 1**. Enriched KEGG pathways for progressive model (PM) and continuous model(CM) genes correspondent to Figure 2F.

**Supplementary file 1: Table S1**. Sample information used in our study with raw file lists and the reads number.

**Supplementary file 2: Table S2**. The number of stages associated gene for each species

**Supplementary file 5: Table S3. Bonferroni-corrected p-values of overlapping CM genes between pairwise species**

**Supplementary file 7: Table S4**. Annotations and GO terms of overlapped genes between CM and CL9.1

**Supplementary file 8: Table S5**. Enriched GO functional annotation for developmental expression patterns CL1-6

**Supplementary file 10: Table S6**. Summary of expressed gene number for each species

